# Developmental heterogeneity of embryonic neuroendocrine chromaffin cells and their maturation dynamics

**DOI:** 10.1101/2022.05.26.493613

**Authors:** Natalia Akkuratova, Louis Faure, Polina Kameneva, Maria Eleni Kastriti, Igor Adameyko

**Author notes:** equally contributing corresponding authors Please address correspondence to or. equally contributing co-first authors (Natalia Akkuratova, Louis Faure).

## Abstract

During embryonic development, nerve-associated Schwann cell precursors (SCPs) give rise to chromaffin cells of the adrenal gland via the “bridge” transient stage, according to recent functional experiments and single cell transcriptomics data from humans and mice. However, currently existing data do not resolve the finest heterogeneity of developing chromaffin populations. Here we took advantage of deep SmartSeq2 transcriptomics sequencing to expand our collection of individual cells from developing murine sympatho-adrenal anlage and uncover the microheterogeneity of embryonic chromaffin cells and corresponding developmental paths. After improving our atlas of sympatho-adrenal development and performing experimental validations, we discovered that SCPs in the local nerve show high degree of microheterogeneity corresponding to early biases towards either Schwann or chromaffin terminal fates. Furthermore, we found that a post-”bridge” population of developing chromaffin cells gives rise to persisting immature chromaffin cells and the two terminal populations (adrenergic and noradrenergic) via diverging differentiation paths. Taken together, we provide a thorough identification of novel markers of adrenergic and noradrenergic populations in developing adrenal glands and report novel differentiation micro-paths leading to them.

## Introduction

Adrenaline and noradrenaline are key catecholaminergic hormones driving stress response in combination with other neurotransmitters as part of the hypothalamus-pituitary-adrenal gland axis (Kvetnansky, Sabban et al. 2009, Eiden 2013). During embryonic development and early postnatal stages, adrenaline and noradrenaline are additionally produced by chromaffin cells of the Zuckerkandl organ (Kohn, Zuckerkandl 1901, Coupland 1965, Böck 1982).

The Zuckerkandl organ is a transient, embryonic chromaffin organ located next to the dorsal aorta and positioned between the kidneys (Ehrhart-Bornstein et al., 2010; Schober et al., 2013). Apart from the different anatomical location of the adrenal medulla and Zuckerkandl organ and the absence of the cortical cells in the later, chromaffin cells that are found in the two organs are considered indiscernible when it comes to their molecular features (Schober, Parlato et al. 2013, Kastriti, Kameneva et al. 2019). In terms of biochemical content, chromaffin cells are classified into two functional subpopulations: the noradrenergic (DBH+/TH+) and adrenergic (DBH+/TH+/PNMT+) cells (Abel 1897, Von Euler 1946, Huber 2006). Potentially, chromaffin cells are even more heterogeneous, as hinted by some previous studies (Shaw and Letourneau 1986, Ohmori 1998, Bedoya-Reina, Li et al. 2021). The previous single cell RNA sequencing of mouse and human adrenals did not reveal much of the unanticipated heterogeneity of chromaffin cells (Furlan, Dyachuk et al. 2017, Dong, Yang et al. 2020, Hanemaaijer, Margaritis et al. 2021, Kameneva, Artemov et al. 2021). However, this might be due to insufficient sampling of chromaffin cells throughout developmental and postnatal stages or shallow recovery of individual transcriptomes. To clarify the situation with minor chromaffin subpopulations, transcriptomic sequencing of higher depth, such as the SmartSeq2 and SmartSeq3 methods, are ideal (Picelli, Bjorklund et al. 2013, Ziegenhain, Vieth et al. 2017, Hagemann-Jensen, Ziegenhain et al. 2020).

As mentioned above, the main function of the adrenal medulla is the preparation of the body for the “fight-or-flight” response through the release of catecholamines. Fetal-derived catecholamines are essential for survival of the fetus during birth and normal initiation of breathing (Lagercrantz and Slotkin 1986, Hillman, Kallapur et al. 2012). Furthermore, secretion of adrenaline by chromaffin cells increases heart rate thus ensuring oxygen delivery to all organs (Abel 1897). Overall, chromaffin cell response to hypoxia includes increased catecholamine secretion, membrane depolarization, voltage-gated Ca^2+^ and inhibition of voltage-dependent K^+^ currents (Nurse et al., 2018; Phillips et al., 2001; Ream et al., 2008; Thompson et al., 2002).

During embryonic development, chromaffin cells are derived from nerve-associated progenitors, or Schwann cell precursors (SCPs), that are derivatives of the neural crest (known as the 4^th^ germ layer) (Furlan, Dyachuk et al. 2017, Lumb, Tata et al. 2018, Kastriti, Kameneva et al. 2019). SCPs follow the preganglionic motor nerves all the way to the sympathoadrenal primordium, where they differentiate towards chromaffin cells via the transient “bridge” state characterized by the specific expression of *Htr3a* and other markers starting from E12-12.5 stage of the mouse embryonic development (Furlan, Dyachuk et al. 2017, Lumb, Tata et al. 2018). Following a peak of chromaffin differentiation between E12.5 and E14.5 stages, the remaining SCPs stay on the nerves as they choose between myelinating and non-myelinating Schwann cell fates. However, it is not clear which molecular events in a subpopulation of SCPs drive the decision-making toward chromaffin fate or mature Schwann cells.

To address this and other questions pertaining microheterogeneity of developing chromaffin cell population, we employed deep single cell RNA sequencing approach SmartSeq2 combined with the trajectory and RNA velocity analysis. Our aim was to visualize the dynamics of the developing adrenal medulla during the generation of chromaffin cells from SCPs and the subsequent generation of an ever-increasing pool of chromaffin cells through self-renewal. In our study, we uncovered and validated previously unknown intermediate chromaffin cell subpopulations and obtained the insights into dynamic heterogeneity among SCPs on the local nerves, which leads to a balanced generation of chromaffin cells and mature Schwann cells necessary for innervation maintenance in the adrenal gland. The new atlas of chromaffin and Schwann cell development is made available online (https://adameykolab.srv.meduniwien.ac.at/ChC/) and will help to address different questions about the development of chromaffin cells, as well as some which might be related to identification of developmental gene expression modules in sympatho-adrenal tumors, such as neuroblastoma and pheochromocytoma.

## Results

### Microheterogeneity unveiled in the nerve-associated cells of the developing sympathoadrenal anlage

In order to systematically analyze the fine aspects of developmental transitions during the differentiation of nerve-associated SCPs towards chromaffin cells and to reveal arising microheterogeneity of neuroendocrine cells, we performed single-cell RNA sequencing using the SmartSeq2 technology of cells of the sympathoadrenal anlage from *Wnt1-Cre;R26R*^*Tomato*^ mice at developmental stages E12.5, E13.5, E14.5, E16.5, E18.5 and P2 (Fig 1A) (Picelli et al., 2013). Although being more expensive and somewhat laborious, SmartSeq2 allows better capture of mRNA transcripts from individual cells as compared to average Chromium 10X results, delivering the average of around 7000 expressed genes from a single cell. Our previous analysis of sympatho-adrenal development with SmartSeq2 resulted in a dataset, which enabled general identification of major transition from SCPs into “bridge” transitory stage and then into chromaffin cells (Furlan, Dyachuk et al. 2017). However, those data did not allow for a deep analysis of chromaffin cell heterogeneity and for the understanding of the mechanisms governing the fate selection process within the nerve towards either mature Schwann or chromaffin phenotypes. In this manuscript, we added more SmartSeq2-sequenced cells from the developing sympatho-adrenal anlage, which resulted in the collection of 1361 single-cell transcriptomes at high sequencing depth, which significantly increased the informative power of our analysis as compared to our previous study (Furlan, Dyachuk et al. 2017).

**Fig.1.**
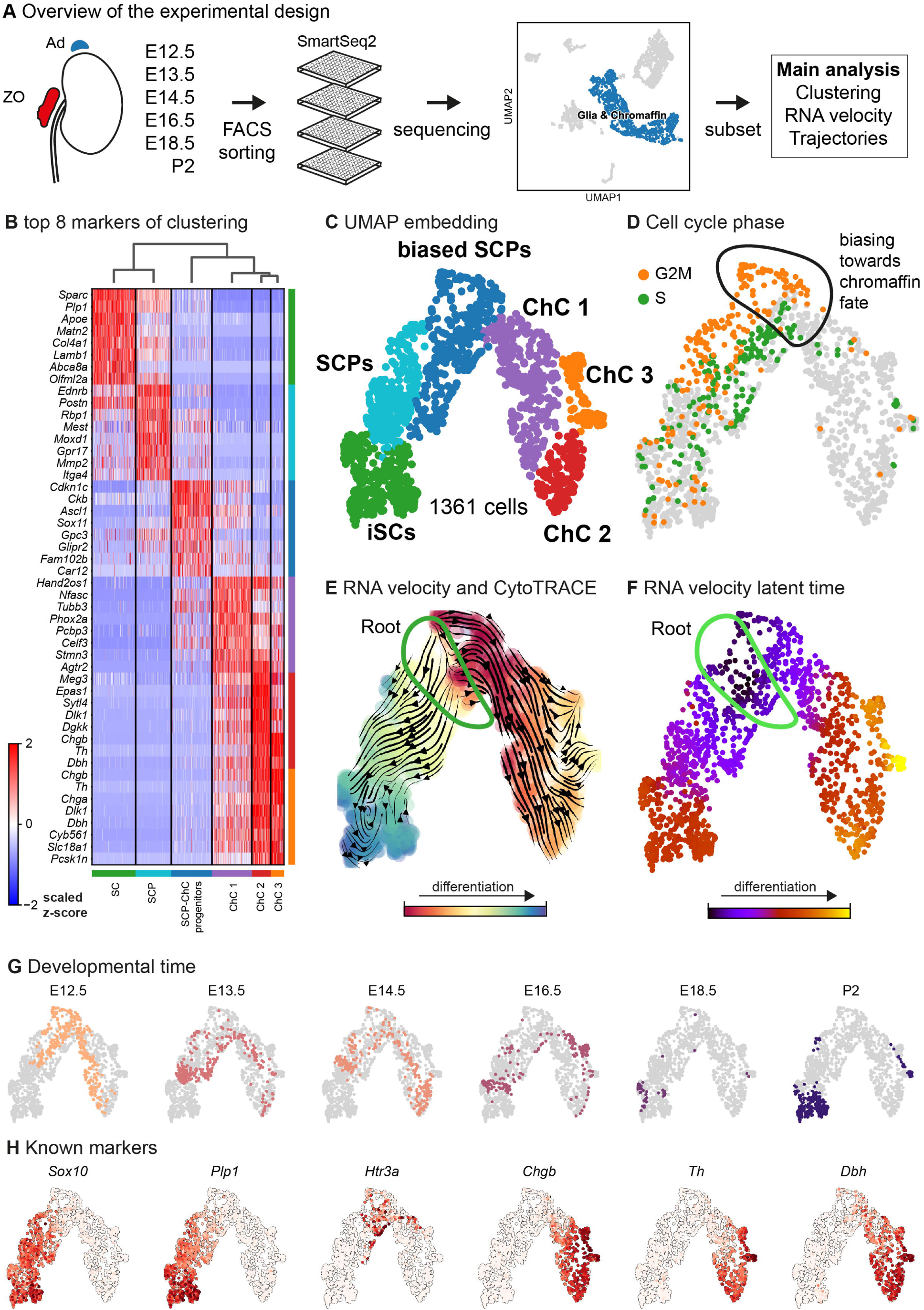
Transcriptomic analysis of the adrenal medulla and Zuckerkandl organ reveals the parallel differentiation of Schwann cell precursors towards Schwann cells and chromaffin cells. ***A*** Overview of the experimental design for the acquisition of *Wnt1*^*TOMATO*^ single-cell transcriptomes of cells of the adrenal medulla and Zuckerkandl organ from stages from E12.5 up to P2, using the SmartSeq2 platform, ***B*** Heatmap showing the top 8 markers of each cluster resulting from the hierarchical clustering of cells with glial and chromaffin markers, ***C*** UMAP embedding of the glial and chromaffin cells from the adrenal medulla and Zuckerkandl organ, ***D*** Cell cycle phase of the selected cells from the adrenal medulla and Zuckerkandl organ, with the first chromaffin fate-biased cells circled, ***E*** Dynamic representation of differentiation dynamics using combined RNA velocity and CytoTRACE with the progenitor Schwann cell progenitor population (SCP) annotated as the root, ***F*** RNA velocity latent time with the progenitor Schwann cell progenitor population (SCP) annotated as the root, ***G*** Developmental time of the single cells shown on the UMAP embedding, ***H*** UMAP of classical markers of Schwann cell precursors and Schwann cells (*So×10, Plp1*), intermediate chromaffin fate-biased “bridge” cells (*Htr3a*) and chromaffin cells (*Chgb, Th, Dbh*) reflecting the main cell types in the data set.

As expected, the primary computational analysis confirmed the presence of Schwann cells precursors (SCPs) (identified by the expression of *Plp1, Ednrb, Itga4* among others) transiting towards either *Plp1+/Sparc+/Postn+* immature Schwann cells or *Th+/Dbh+* chromaffin cells, through the previously described transient “bridge” state characterized by *Htr3a* expression (Fig. 1B,C,H) (Furlan, Dyachuk et al. 2017).

Deeper analysis of the “root” population of SCPs showed the presence of fate-driving biases towards or Schwann cell terminal fate (expressing early known markers *Ngfr, Foxd3, Ednrb, Mpz*) (Fig 2A, B) or sympatho-adrenal (expressing markers *Elavl3, Isl1, Hand2* and *Phox2a*) (Fig 2C,D), which are defined via expression of early fate-related gene modules with a gradual onset of expression (Soldatov, Kaucka et al. 2019). The antagonizing genes expression programs co-exist in cells of the “root” SCP population (Fig 3A-C), which corresponds to the previously described mechanism of fate selection including early co-activation of competing gene expression programs (Soldatov, Kaucka et al. 2019). These biases arise in a “root” SCP population that will transit towards the immature Schwann cell fate (as seen by early *So×10, Plp1, Foxd3* expression and late *Mpz* expression onset) and progress fast along the trajectory towards terminal states (Fig 3B, Fig 2B,C). This process coincides with the gradual decrease of a proportion of proliferative differentiating cells (Fig 1D). The expression of mature Schwann-cell related genes emerges next to the cells primed for sympatho-adrenal differentiation or even in the same cells. The priming of “root” SCPs towards different fates suggests a significant degree of microheterogeneity in nerve-associated cells of developing sympatho-adrenal anlage.

**Fig.2.**
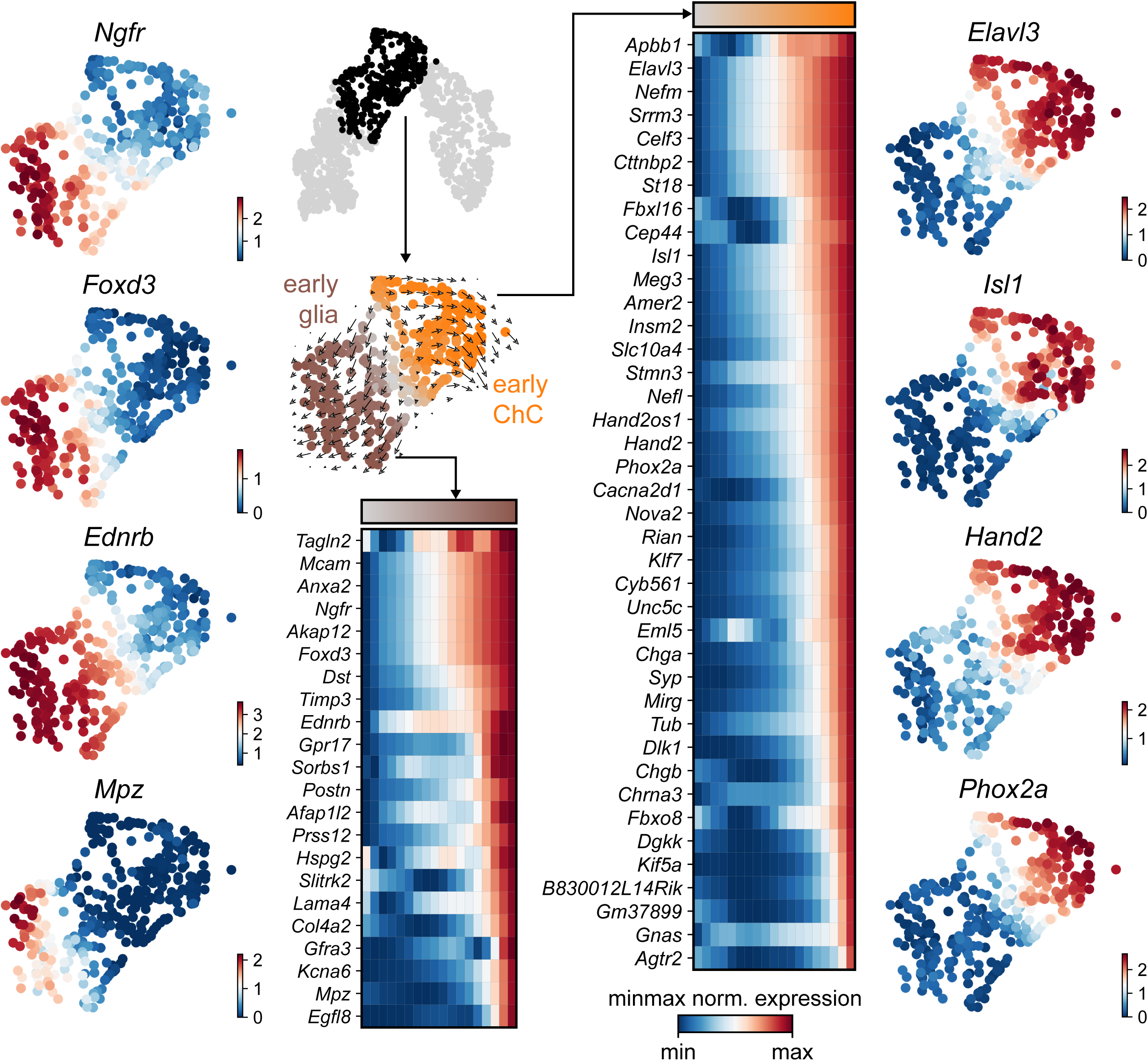
Trajectory analysis from SCP diverging to glial and ChC fates. ***A*** Selected early diverging glial markers. ***B*** Sub-selection of the diverging pseudotime trajectory (Furlan, Dyachuk et al.), confirmed by RNA velocity (middle). Binned (10 bins) pseudotime heatmap of min-max normalised expression of significantly changing markers leading to glial fate, ordered by activation (Cox, Sadlon et al.). ***C*** Binned (10 bins) pseudotime heatmap of min-max normalised expression of significantly changing markers leading to ChC fate, ordered by activation. ***D*** Selected early diverging ChC markers.

**Fig. 3.**
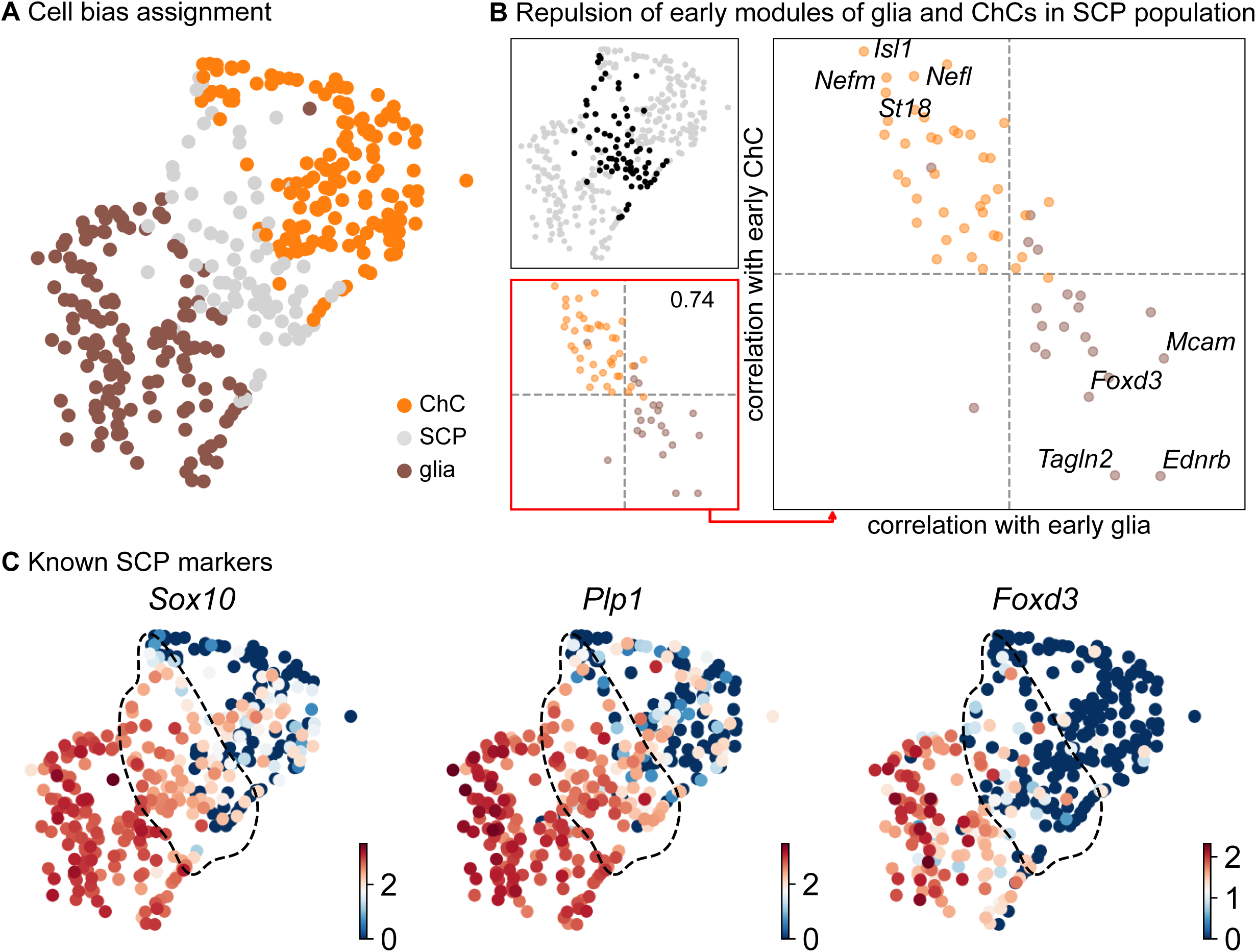
Compositional and biasing analysis of SCP cells toward either of the fates. ***A*** scoring of the cells using the two list of early diverging genes define in Fig. 2, with cells considered SCP when the scoring is low. ***B*** Gene module repulsion analysis between early diverging glia and ChC gene groups. Analysis is performed only in SCP cells (top left). The scatter plot (lower left and right side) depicts inter and intra-module correlation of each gene from both diverging gene group. A repulsion score is shown in the lower left plot. Top 4 gene showing highest intra-module correlation and inter-module anti-correlation are annotated on the right-side plot. ***C*** UMAP plots of log10(fpm) expression of three known markers, with SCP cells highlighted by the dashed lines.

Contrary to SCPs biased towards Schwann cell fate, SCPs primed towards chromaffin fate start to show the early expression of *Sparc, Birc5* and *Ccnb2* genes, which is followed by the classic “bridge” signature and chromaffin differentiation. The biasing of SCPs towards chromaffin fate coincides with dramatic switch in the mitotic index (Fig 1D) and, as expected, is identified by the expression of proneurogenic genes *Ascl1* and *Sox11* (Fig 1B), followed by gradual upregulation of *Phox2a, Tubb3, Hand2, Agtr2* and others. The corresponding cluster of differentiating cells (ChC1) is composed of cells ranging from E12.5 to E16.5 and corresponding to cells that are not proliferating (Fig 1D,E,F,G), which suggests that multiple waves of chromaffin fate-biased cells are recruited from SCPs residing on the local innervation and maturing towards chromaffin phenotype up to E16.5. This is further confirmed by the observation of gradual upregulation of known chromaffin markers *Th, Dbh* and *Chgb* (Fig 1B,H) and RNA velocity analysis combined with CytoTRACE, as well as latent pseudotime analysis (Fig 1E,F).

### Closely nested paths lead towards different chromaffin end-point states

Further analysis identified the developmental progression of early pro-chromaffin cells of ChC1 cluster towards two endpoints composed of smaller subpopulations of *Th*^*high*^*/Dbh*^*high*^*/Chgb*^*high*^ chromaffin cells arising from E16.5 and P2 timepoints (together described as ChC3) and *Th*^*low*^*/Dbh*^*low*^*/Chgb*^*low*^ cells (ChC2) originating predominantly from earlier E12.5-E14.5 stages with only some cells coming from E16.5 stage (Fig.1C,G). The terminal subpopulation of chromaffin cells of ChC2 cluster is interpreted as a persisting immature chromaffin state that will later become mature and will choose a fate of one of the fully differentiated chromaffin endpoints. The re-clustering of the chromaffin branch and splitting ChC2 and ChC3 into finer clusters in combination with RNA velocity shows that ChC3 cells contain the two terminal subpopulations of adrenergic (*Th*+/*Dbh*+/*Pnmt*+ - cluster 14) and noradrenergic (*Th*+/*Dbh*+/*Pnmt*- - cluster 15) chromaffin cells (Fig 1C,E,F and Fig 4A,B). The adrenergic subpopulation additionally showed a specific enrichment for the expression of *Rab3b* and higher levels of *Chga, Th, Dbh, Scg2* and *Npy*, whereas the noradrenergic subpopulation showed specific expression of *Gm2115, Cd27, Lamc3* and *Hif3a*. The bifurcation process reflected in sub-trajectories leading to these two terminal subpopulations emerge within cluster 8, which also forms a branch towards the endpoint with immature chromaffin ChC2 cells. Cluster 8 belongs to post-”bridge” chromaffin differentiation, which is positive for *Htr1b and Mei4* RNAs and contains microheterogeneous population of cells biased towards the immature chromaffin state of ChC2 cluster or cells transiting into adrenergic and noradrenergic subpopulations. The differentiation paths predicted by RNA Velocity suggest that terminal populations, although taking neighboring positions, are not interconverting or giving rise to each other, and instead require a fate selection process in the progenitors positioned downstream the differentiation trajectory. Thus, post-”bridge” immature chromaffin cells undergo fine fate and state selection processes to yield functional heterogeneity of the chromaffin cells in the developing adrenal medulla.

**Fig.4.**
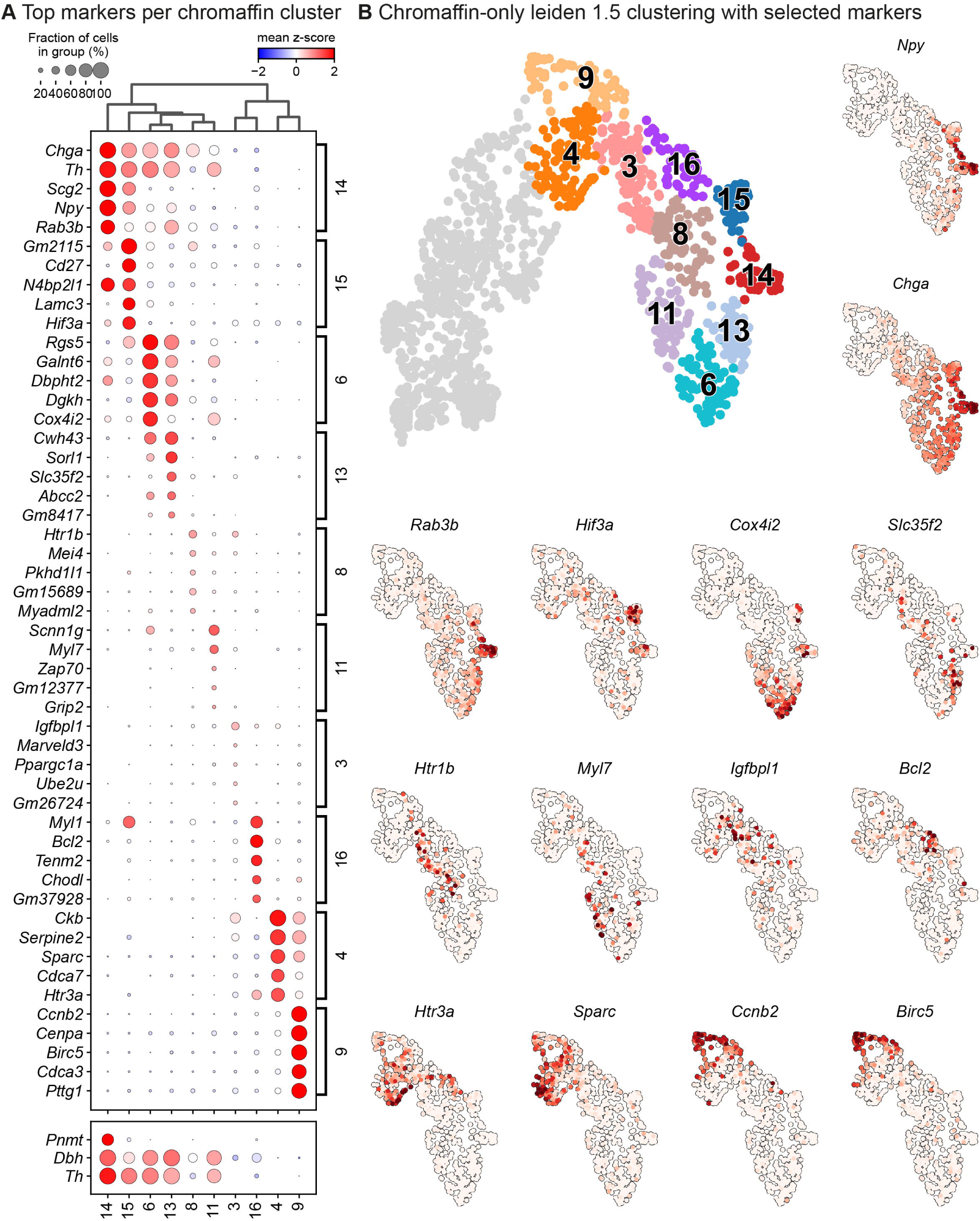
Leiden analysis shows ten main clusters of chromaffin cells based on its gene expression in murine adrenal medulla from E13.5 to P2 developmental stages. ***A*** Dot plot showing top 5 markers identified by gene expression for each leiden cluster of chromaffin cells. ***B*** Unbiased plotting using Leiden clustering presented via UMAPs with highly expressed markers in different clusters.

**Fig.5.**
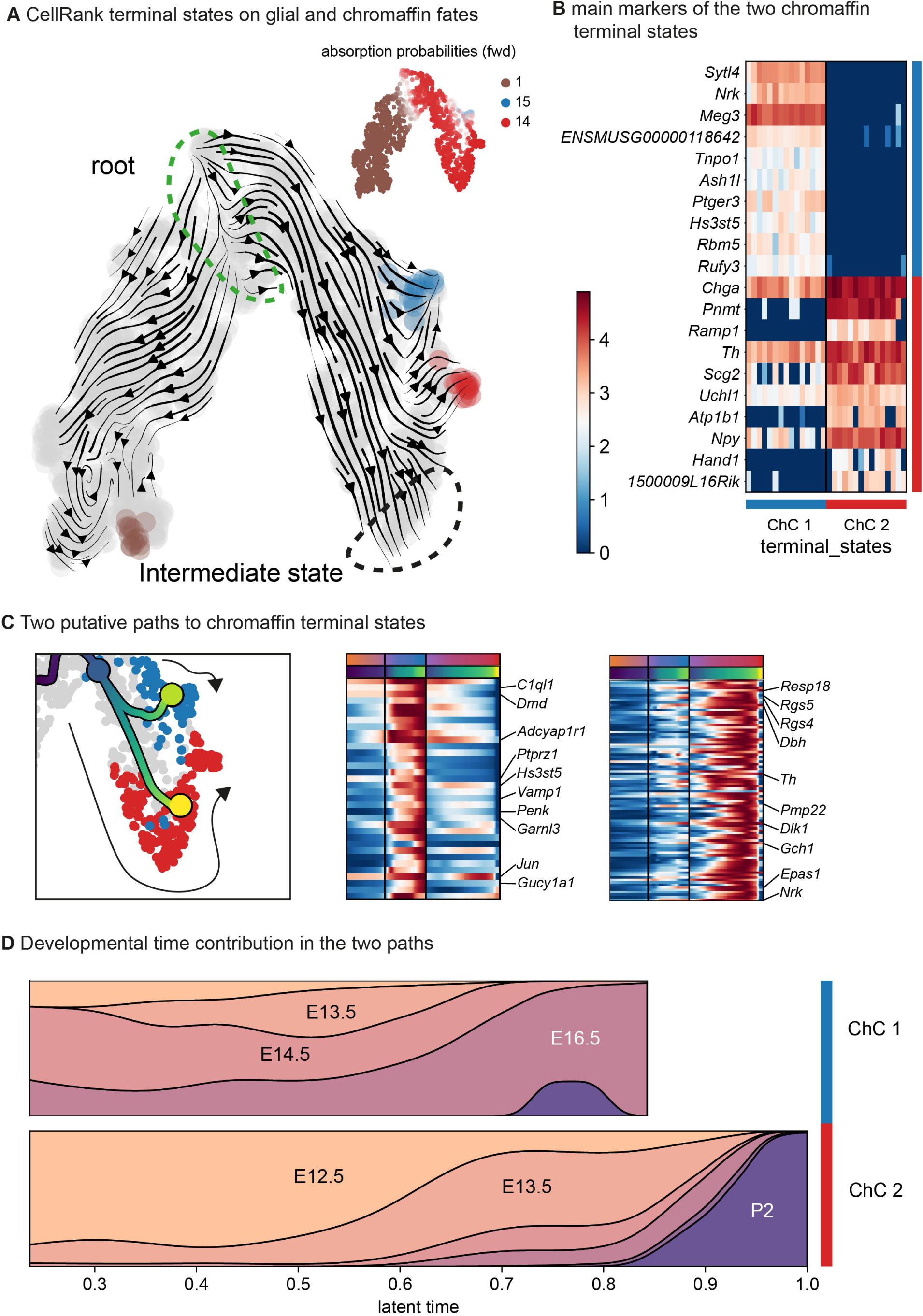
Expression of cluster-specific markers in embryonic murine adrenal medulla at E14.5 and E16.5. ***A, B*** Immunohistochemistry for TH and RNA *in situ* hybridization for *Rgs5* and *Myl1* on cross-sections of embryonic adrenal medulla at E14.5 and E16.5 accordingly. Scale bar is 10 μm, scale bar on insets is 2 μm. ***C, D*** Immunohistochemistry for TH and RNA *in situ* hybridization for *Notum* and *Scg2* on cross-sections of embryonic adrenal medulla at E14.5 and E16.5 accordingly. ***E, F*** Immunohistochemistry for TH and RNA *in situ* hybridization for *Lrp1b* and *Scnn1g* on cross-sections of embryonic adrenal medulla at E14.5 and E16.5 accordingly.

### Novel subpopulations of chromaffin cells correlate with maturation of the gland and functional heterogeneity

The refinement of the general clustering by re-clustering only the branch differentiating towards chromaffin cells, resulted in 6 subpopulations that arise at different time points during embryonic development and two endpoints of mature adrenergic and noradrenergic chromaffin cells (Fig 4A,B – clusters 3, 6, 8, 11, 13, 14 correspond to novel populations, while 15, 16 are mature chromaffin cells, all of which are identified as chromaffin cells based on *Th* and *Dbh* expression). The three larger subpopulations from the general previous clustering of chromaffin cells can be further split into smaller subpopulations based on specific markers: the former ChC1 cluster from more general clustering includes clusters 3, 8, 11, and 16. These clusters reflect the early steps of chromaffin cell development, which include a transient “bridge” (expressing *Htr3a*) population and immediate post-bridge immature chromaffin cells.

Cluster 3 is characterized by downregulation of glial identity, as indicated by the expression of negative regulators of myelination such as *Tmem98* and *Jam2*, as well as *Notch1* and *Sox11*, implicated in negative regulation of glial proliferation. Cluster 8 comprises of cells gaining endocrine identity, by expressing neurotransmitter and synaptic vesicle related markers such as *Cdk5, Pclo, Ank2* and *Dmd*. Cluster 11, while being transcriptionally similar to cluster 8, is characterized by genes related to axon guidance (*Ret, Robo2* and *Sema4f*) and by gene implicated in noradreline biosynthesis (*Th, Dbh* and *Insm1*). Cluster 6 further diverges from all described clusters, with cells defined by *Rgs5* expression, representative of committed chromaffin progenitors. This cluster is further defined by markers related to synaptic transmission and axonal processes, with the expression of *Gria2, Syt1, Npy* and *Kcnc4*.

The general ChC2 population from the previous clustering includes clusters 6 and 13, which, as discussed above, are characterized by high expression of the oxygen-sensing-related genes *Epas1 (*also known as hypoxia-inducible factor, *HIF-2*α) and *Cox4i2* (Wang, Jiang et al. 1995, Yang, Sun et al. 2013, Zhou, Chien et al. 2016) (Fig. 1A, Fig 4B). The most advanced general subpopulation ChC3 consists of clusters 14 and 15 which share the expression of chromaffin markers *Th, Dbh, Chga* and *Npy*. To validate the existence of fine subpopulations of chromaffin cells, we analyzed the developing adrenal glands on stages E14.5, E16.5, E18.5 and P2 by immunohistochemistry and fluorescent RNAscope® *in situ* hybridization using unique markers for one or several subpopulations (Fig.6,7).

**Fig.6.**
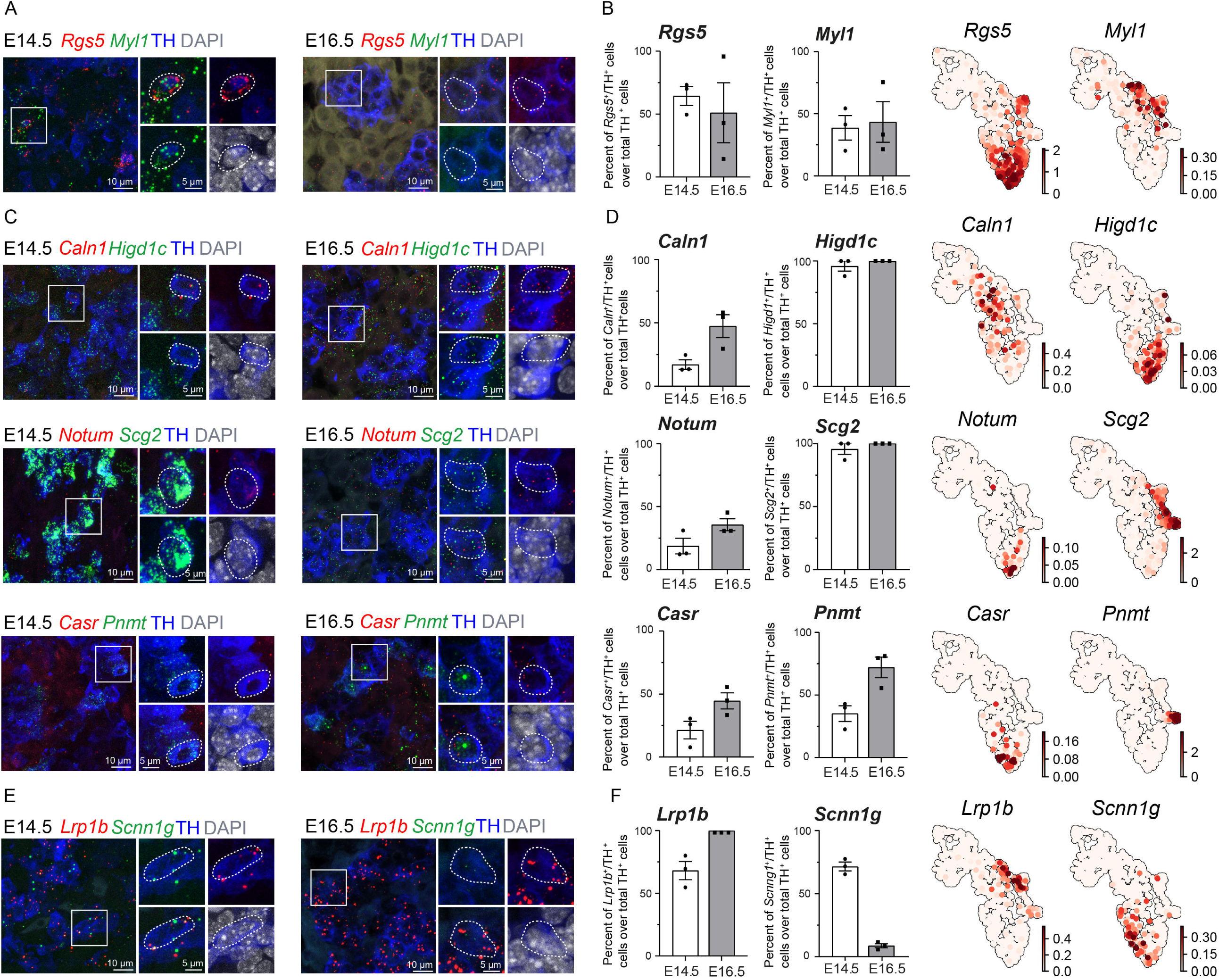
Expression of cluster-specific markers in murine adrenal medulla at E16.5 and P2. ***A, B*** Immunohistochemistry for TH and RNA *in situ* hybridization for *Rgs5* and *Myl1* on cross-sections of embryonic adrenal medulla at E18.5 and P2 accordingly. Scale bar is 10 μm, scale bar on insets is 2 μm. ***C, D*** Immunohistochemistry for TH and RNA *in situ* hybridization for *Notum* and *Scg2* on cross-sections of embryonic adrenal medulla at E18.5 and P2 accordingly. ***E, F*** Immunohistochemistry for TH and RNA *in situ* hybridization for *Lrp1b* and *Scnn1g* on cross-sections of embryonic adrenal medulla at E18.5 and P2 accordingly.

**Fig.7.**
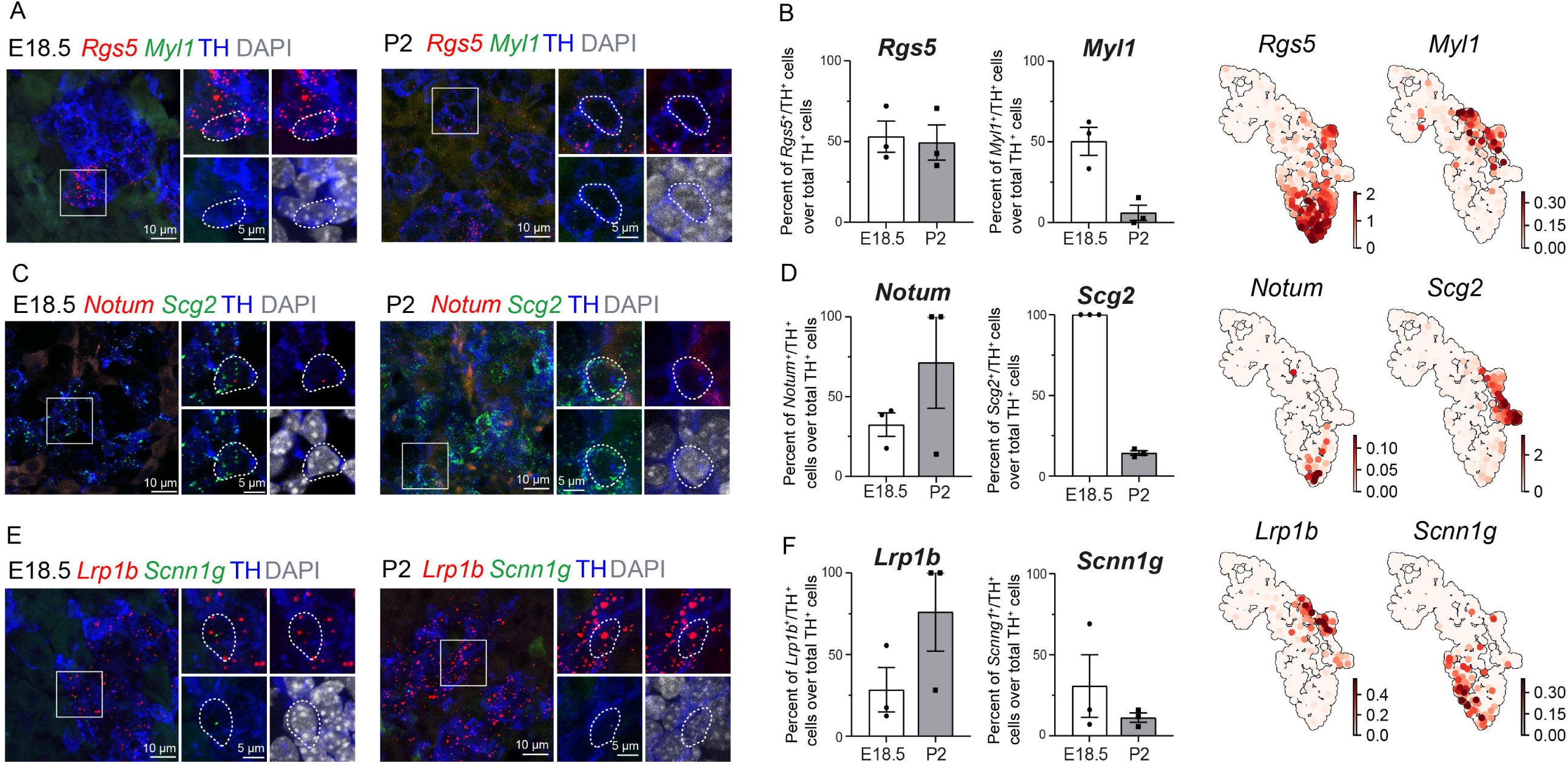
Identification of two discrete paths of chromaffin cell differentiation. ***A*** Dynamic representation of differentiation dynamics using RNA velocity identifies two terminal states of chromaffin fates (red and blue) with the Schwann cell progenitor population (SCP) annotated as the root. ***B*** Heatmap presents main markers of the two chromaffin terminal states. ***C* Left panel:** Two putative paths to chromaffin terminal states. Middle and **right panels:** Heatmaps showing unbiased markers for each chromaffin terminal state. ***E*** Developmental time contribution of chromaffin cells to the two terminal paths.

According to scRNA-seq, *Rgs5* is specifically expressed in clusters 15, 6 and 13 (Fig. 4A). While clusters 6 and 13 belong to the oxygen-sensing type of chromaffin cells, cluster 15 has a unique expression of *Myl1*. Therefore, by combination of these markers with the pan-chromaffin cell marker TH (tyrosine hydroxylase) we show that the proportion of *Rgs5*^+^ cells ranging from 64.30±7.44 to 49.41±10.80 % during stages 14.5-P2 (Fig. 6A, B, 7A, B). The expression of *Myl1* is at its widest at E18.5 with 50.29±8.72% of chromaffin cells showing high levels of expression (Fig. 6B). Post-natal expression of the gene drops dramatically with minimal expression of *Myl1* in 5.99±4.76% of cells at P2 (Fig.6A, B, 7A, B). This population almost completely seizes at P2, rendering its role the most important during embryonic development of adrenal medulla, but not at postnatal life.

*Rgs5* gene, coding for a regulator of G-protein signaling 5, a marker that is involved in development of the adrenal medulla, is expressed at approximately similar level at all developmental stages. It has been also shown as a hypoxia-inducible apoptotic stimulator in endothelial cells (Jin, An et al. 2009, Chan, Komada et al. 2019). *Caln1* gene, coding for Calcium-binding protein 8, was shown previously only in adrenal cortex as a main regulator of Ca^2+^ storage but in our study is found in all subpopulations of chromaffin cells from early developmental stages (Kobuke, Oki et al. 2018). The only exception is cluster 14, which corresponds to adrenergic chromaffin cells. *Higdc* gene, coding for Hypoxia Inducible Domain Family Member 1C, is also expressed in all subpopulations except 3 at E12.5-E16.5. It was shown to be highly expressed in carotid bodies in comparison with adrenal medulla (AM) (Chang, Ortega et al. 2015). Though AM realises catecholamines in response to changes of oxygen level in blood, carotid bodies is the major oxygen-sensing organ in human body (Chang, Ortega et al. 2015). *Myl1*, coding for Myosin Light Chain 1, was previously shown only in muscle tissue and endocrine cells of pituitary gland. Here we show that it is expressed mostly in late developmental stages E16.5-P2, in clusters 15 and 16 (Welcker, Hernandez-Miranda et al. 2013). *Notum*, which encodes Palmitoleoyl-Protein Carboxylesterase, is one of unique markers that expresses mostly in subpopulations 6 and 13. We have shown that its expression rises with the development of AM and is highest at postnatal stage P2. *Notum* has been shown to inhibit WNT signalling and is required for vertebrate brain development (Kakugawa, Langton et al. 2015, Zhang, Cheong et al. 2015) *Scg2*, encodes secretogranin II, that is responsible for packing and sorting neurotransmitters in secretory vesicles in adrenal gland, and has highest levels of expression in subpopulations 14 and 15 (endpoints) (Steiner, Schmid et al. 1989, Fang, Dai et al. 2021). *Scnn1g*, which encodes Sodium Channel Epithelial 1 Subunit Gamma, is expressed in all subpopulations except 3 (Fig. 6,7).

Another important marker is *Lrp1b*, Low-density lipoprotein receptor-related protein 1B, as mutations in this gene were shown to play a role in the development of paraganglioma and pheochromocytoma (Dong, Huang et al. 2020, Kudryavtseva, Kalinin et al. 2020, Choi, Lim et al. 2021). We analysed the heterogeneous expression of *Lrp1b* and found it expressed specifically in noradrenergic mature chromaffin cells, as expected from the computational analysis (Fig 6,7). The time-course of *Lrp1b* expression suggested its dynamic expression with specific role during development.

Lastly, to uncover underlying maturation dynamics we examined separately all chromaffin clusters that are not assigned to the two endpoints of noradrenergic (cluster 15) and adrenergic (cluster 14) or composed of “bridge” cells (Fig 8). The result of this analysis on the subselected clusters 3, 6, 8, and 11 revealed a progressive upregulation of chromaffin markers *Dbh* and *Th* with cluster 3 characterized with lowest levels and cluster 6 manifesting the maximum expression (Fig 8A). Additionally, each cluster was characterized by the expression of specific markers, as well as the progressive upregulation from one to the next, resembling a pseudotime progression (Fig 8B, C).

**Fig.8.**
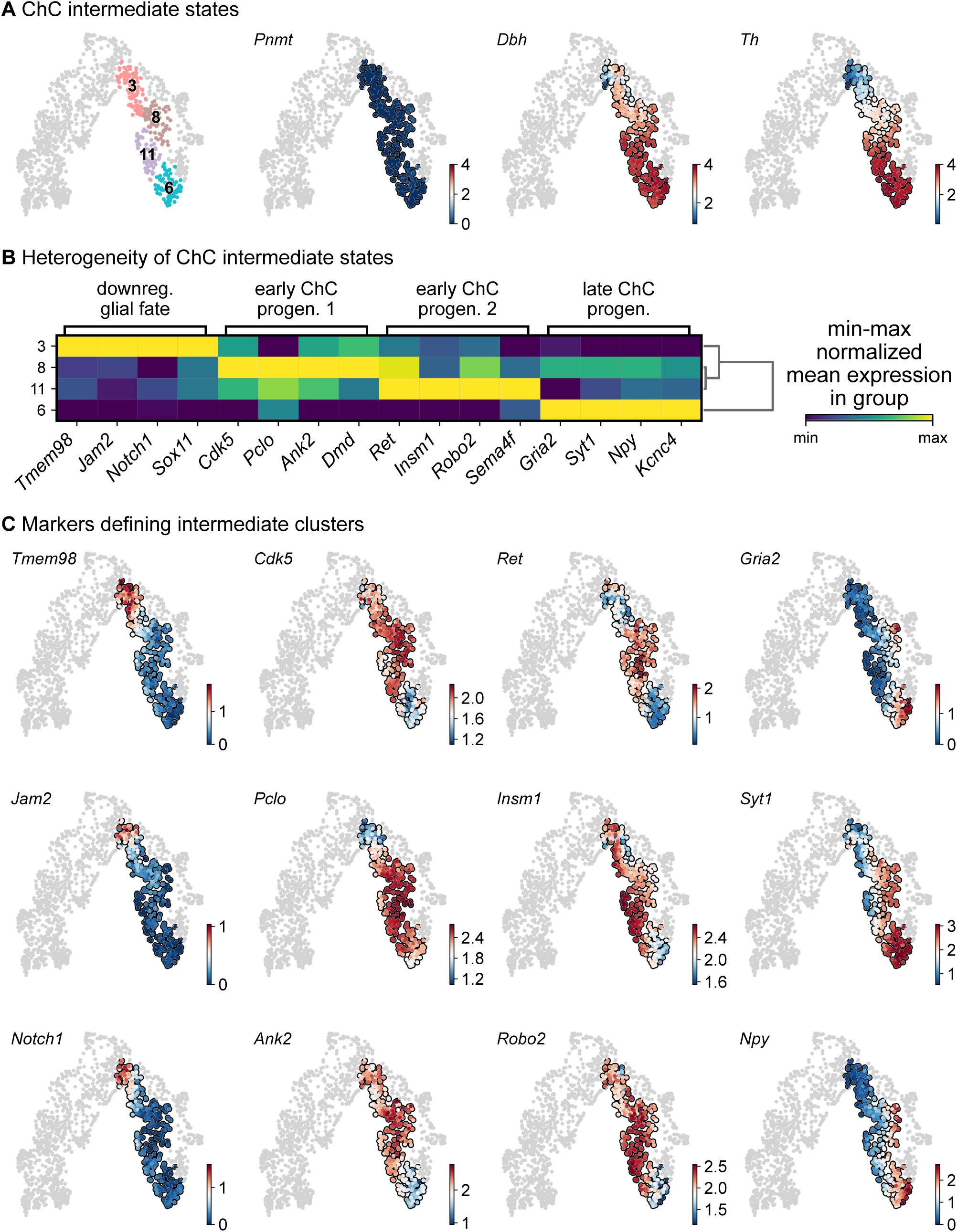
Heterogeneity of intermediate states of chromaffin cells. ***A*** Selection of intermediate leiden clusters shown on UMAP, with three known markers represented in that subselection (knn smoothed log10 fpm). ***B*** Matrix plot of min-max normalized mean expression of selected DE markers from each leiden clusters. ***C*** UMAP of the selection markers for each leiden clusters (knn smoothed log10 fpm).

Overall, this analysis of single cell RNAseq data and microheterogeneity of developing mouse adrenal gland predicted previously unrecognized fine transitory steps already after the “bridge” state, which lead to fate partitioning towards two maturing and one immature persisting population of chromaffin cells. The experimental validations confirmed the structure of predicted transient and terminal populations.

## Discussion

The recent successes in single cell transcriptomics helped to resolve the heterogeneity of developing adrenal medulla, focused on major developmental transitions from nerve-associated SCPs to then-unknown “bridge” cells and to chromaffin cells (Furlan, Dyachuk et al. 2017, Dong, Yang et al. 2020, Bedoya-Reina, Li et al. 2021, Hanemaaijer, Margaritis et al. 2021, Kameneva, Artemov et al. 2021). Although the general developmental trajectory of sympatho-adrenal cells created a lot of new understanding of the processes in developing sympatho-adrenal anlage, fine heterogeneity of nerve-associated SCPs or chromaffin cells stayed rather enigmatic, because single cell transcriptomics is sensitive to insufficient sampling and sequencing depth (Ziegenhain, Vieth et al. 2017). In this study, we generated deep single cell transcriptomic data from all stages of chromaffin development, including early postnatal stages, aiming to identify populations that were previously invisible due to lower sampling and incomplete coverage of developmental timing. We utilized SmartSeq2, which allows the recovery 7000 - 8000 expressed genes per cell on average to provide the necessary depth (Picelli, Bjorklund et al. 2013, Ziegenhain, Vieth et al. 2017). The analysis of this atlas showed that we moved beyond the previous understanding of sympatho-adrenal development and identified heterogeneity of SCPs in the nerve and novel post-“bridge” fate splits leading to previously unrecognized heterogeneous populations of chromaffin cells.

The SCPs, neural crest-derived nerve-associated stem cells (Furlan and Adameyko 2018), cover the nerve surfaces and travel with the nerves to almost all locations in the developing embryo. In some of the locations, they detach and differentiate into a number of tissue-specific cells types, such as melanocytes, parasympathetic and sympathetic neurons, enteric neurons, chromaffin cells, oxygen-sensing glomus cells and specific mesenchymal cells (Joseph, Mukouyama et al. 2004, Adameyko, Lallemend et al. 2009, Nitzan, Pfaltzgraff et al. 2013, Dyachuk, Furlan et al. 2014, Kaukua, Shahidi et al. 2014, Uesaka, Nagashimada et al. 2015, Espinosa-Medina, Jevans et al. 2017, Furlan, Dyachuk et al. 2017, Hockman, Adameyko et al. 2018, Kastriti, Kameneva et al. 2019). The multipotency of SCPs makes them similar to their maternal population – the migratory neural crest, often called the 4^th^ germ layer (Bronner and Simoes-Costa 2016, Furlan and Adameyko 2018). During the development of adrenal medulla, SCPs covering the motor axons extending from neural tube, become recruited during adrenal gland development and differentiate into chromaffin cells and intra-adrenal sympathoblasts (Furlan, Dyachuk et al. 2017, Lumb, Tata et al. 2018, Kameneva, Artemov et al. 2021). This process was previously dissected using single cell transcriptomics and lineage tracing (Furlan, Dyachuk et al. 2017, Kameneva, Artemov et al. 2021), and yet, we still do not understand why some SCPs become recruited towards sympathoadrenal differentiation and detach from the nerve while others remain on the nerve and differentiate towards mature Schwann cells. Here we attempted to address the underlying molecular mechanisms by analyzing the divergence of the differentiation directions within a SCP population, which surprisingly suggested high microheterogeneity already within the nerve-residing population. This microheterogeneity of SCPs was related to the expression of transcriptional programs related to either chromaffin or Schwann cell fates and included *Ascl1, Hand2, Phox2a/b, Insm2* genes (chromaffin fate) or *Ednrb, Foxd3, Postn, Itga4, Ngfr* and *Gfra3* genes (Schwann cell fate). The discovered fate-related biases (emerging as coordinated transcriptional programs) within the population of nerve-associated cells suggest that spinal motor neurons and their axons might play a role in providing signals stabilizing the SCP and Schwann cell phenotype, also by antagonizing the signals from the local tissue and inhibiting the recruitment of SCPs, which drives the detachment from the nerve. One of such nerve-borne stabilizing signal might be NRG1, which is a well-known factor maintaining the survival and phenotype of SCPs (Britsch, Li et al. 1998, Taveggia, Zanazzi et al. 2005). We anticipate that our single cell data on the biased SCP population will help to advance the understanding of biasing extrinsic signals, and will provide educated guesses for future functional experiments.

Cell type heterogeneity of chromaffin cell population was studied and discussed for a long period of time (Wurtman, Axelrod et al. 1968, Bohn, Goldstein et al. 1981, Bedoya-Reina, Li et al. 2021). The classic knowledge stipulates that there are two major subpopulations of chromaffin cells. One of those populations synthesizes noradrenaline, and the other one is responsible for the synthesis of adrenaline. Although both populations express TH and DBH, only the second population has an additional enzyme PNMT, which converts noradrenaline to adrenaline (Von Euler 1946, Huber 2006). However, a recent study shed new light on chromaffin heterogeneity, reporting several chromaffin subtypes (Bedoya-Reina, Li et al. 2021). In our study, we revealed 8 subpopulations of developing chromaffin cells with distinct temporal dynamics and maturation endpoints. Whereas most of these populations appeared transient, three of them turned out to be terminal states. Two of those states reflected the adrenergic and noradrenergic chromaffin cells while the third one was characterized by robust expression of genes associated with oxygen-sensing.

Specifically, chromaffin cells take part in response to hypoxia mediated by increased catecholamine secretion (Abel 1897, Von Euler 1946, Coupland 1965, Böck 1982, Kvetnansky, Sabban et al. 2009). Similar to chromaffin cell of adrenal gland medulla and organ of Zuckerkandl, catecholaminergic glomus cells (one of the two cell types, which compose carotid bodies) are also oxygen-sensing and show the presence of a similar transcriptional signature (Zhou, Chien et al. 2016) as well as similar embryonic genesis, which includes contribution from glial nerve associate progenitors (Hockman, Adameyko et al. 2018). The major genes known to be involved in oxygen sensing in adrenal gland include *Epas1* (or *Hif2a*) and *Cox4i2* (Yang, Sun et al. 2013, Zhou, Chien et al. 2016) and are indeed enriched in some of the subpopulations (mostly subclusters 6, 11 and 13, making up ChC2).

Some of the terminal population-specific markers, for instance *Higd1c, Rgs5 and Scg2* are involved in the downstream response following hypoxia-sensing (Jin, An et al. 2009, Fang, Dai et al. 2021, Timón-Gómez 2021). We experimentally validated their expression in adrenal medulla and found population-specific expression in the previously identified oxygen-sensing cells (cluster ChC2). It is plausible that chromaffin cells of the corresponding populations are involved in different and yet specific mechanisms of the oxygen sensing and might show individual features of their transcriptional response to hypoxia. This direction of work will require functional experiments and precise characterization of the differences in hypoxia sensing mechanisms in different chromaffin cells.

The transitions towards terminal chromaffin populations showed a previously unrecognized fate-split in the post-”bridge” population of immature chromaffin cells. Therefore, some immature chromaffin cells represent a pool of undecided progenitors that can balance the proportions of terminally differentiated chromaffin populations during the development and beyond.

As tumor cells re-play the developmental gene expression programs to metastasize, exert phenotypic plasticity and are often resistant to treatment, it is of interest to have a proper description of fine developmental transitions and gene expression modules related to healthy cell phenotypes, developing and terminally differentiating. For instance, this might be relevant for better understanding of intra- and inter-tumor cell heterogeneity of pheochromocytoma and paraganglioma, which are neuroendocrine tumors that originate from chromaffin cells or their progenitors (Mete, Asa et al. 2022). In our study, we took into the account the role of some genes in these tumors, and, thus, selected the markers for experimental validations of identified chromaffin subpopulations keeping such genes in mind. For instance, one of such subpopulation-specific markers is *Lrp1b*, and the mutations in *Lrp1b* lead to development of malignant pheochromocytoma and paraganglioma (Dong, Huang et al. 2020, Kudryavtseva, Kalinin et al. 2020, Choi, Lim et al. 2021). *Rgs5* is another selected marker, which is specific for only some of the identified populations, and has a high level of expression in pheochromocytoma (Choi et al., 2021; Rouillard et al., 2016).

Overall, we provide the detailed single cell transcriptomic atlas of chromaffin cells development, which allows identification of previously unrecognized populations and helps to establish fine transitions in the populations of immature chromaffin cells. Using this updated atlas, we identified and characterized emergence of subpopulations of chromaffin cells that were not known from before, described their potential role in oxygen-sensing and addressed microheterogeneity of the nerve-associated SCPs giving rise to chromaffin and Schwann cells during development. Being easily browsable and accessible online, the extended atlas of Schwann cell and chromaffin cell development will help to advance the field of sympatho-adrenal development, as well as it might be useful for neuroblastoma and pheochromocytoma research communities for better understanding of tumor cell type heterogeneity by comparing malignant cells with cell states in normal chromaffin cell development.

## Material and methods

### Animals

*Wnt1-Cre;R26R*^*Tomato*^ mice (*Wnt1-Cre;R26R*^*Tomato+*^ for single cell preparation and *Wnt1-Cre;R26R*^*Tomato-*^ littermates for the biological validations) were used in all experiments. Stocks were obtained from the Jackson Laboratory: *Wnt1-Cre* (stock #003829) and *R26R-Tomato* mice (stock #007914). The day of plug detection was considered as E0.5 and the day of delivery as P0. 2-6 month old animals were used for mating; the embryos E12.5, E13.5, E14.5, E16.5, E18.5 and postnatal animals P2 of both genders were used in the study. All experiments were performed under the permission of the Jordbruksverket De regionala Djurförsöksetiska nämnderna and Stockholms Djurförsöksetiska nämnd, permit number Igor Adameyko: N15907-2019 and according to The Swedish Animal Agency’s Provisions and Guidelines for Animal Experimentation recommendations.

### Single cell preparation for transcriptomics

Isolated adrenal glands and organ of Zuckerkandl were separately transferred on a plate with HBSS and cut into small pieces using a clean scalpel. Dissociation was performed using 0,05% Trypsin/0,02% EDTA for 15 minutes at 37°C and triturated with P-1000 and P-200 pipettes tips until complete tissue dissociation. Following the addition of equal volume of 10% FBS, the tissue suspension was centrifugated at 500 g, 5 min, 4°C. Following three washes with 10% FBS the pellet was resuspended in 1xPBS. Cells were sorted using a BD FACSAria III cell sorter into 384-well-plates and stored at −80°C until sequencing took place according to a published protocol (Picelli, Bjorklund et al. 2013).

### Generation of count matrices, QC and filtering

The single-cell transcriptome data were generated at the Eukaryotic Single-cell Genomics facility at Science for Life Laboratory in Stockholm, Sweden. The samples were analysed by first demultiplexing the fastq files using deindexer (https://github.com/ws6/deindexer) using the nextera index adapters and the 384-well plate layout. Individual fastq files were then mapped to GRCm39 vM27 genome with ERCC annotations using the STAR aligner using 2-pass alignment (Dobin, Davis et al. 2013). Reads were filtered for only uniquely mapped and were saved in BAM file format; count matrices were subsequently produced using featureCounts. Estimated count matrices were gathered and converted to an anndata object. Cells having more that 5×10^4^ transcripts, 3000 detected genes or less than 25% of ERCC reads were kept. The resulting filtered count matrix contained 2760 high quality cells from all developmental stages.

### Initial clustering with scanpy and scFates

The initial analysis of the count matrix was performed using the scanpy and scFates python packages (https://github.com/kharchenkolab/pagoda2 and https://github.com/LouisFaure/scFates). Overdispersed genes were detected using pagoda2 approach (scFates, pp.find_overdispersed). PCA was performed on the scaled overdispersed genes (scanpy, pp.pca, default parameters) and used for nearest neighbors graph generation (scanpy, pp.neighbors, k=15, Euclidean distance) and UMAP visualisation (scanpy, tl.umap,). (Becht et al., 2019) Clustering were performed from the generated neighbor graph, with a low resolution to obtain a general cell-type overview (scanpy, tl.leiden resolution=0.1) (Traag et al., 2019) Clusters containing cells expressing *So×10, Isl1 or Chga* were included in the downstream analysis. The subset count matrix was further processed similarly, except that cell cycle genes were removed from the highly variable genes. Leiden clustering was performed at resolutions of 0.3 for overview and 1.5 for precise mapping. Cell-cycle was scored using scvelo function scv.tl.score_cell_cycle. Finally, the raw counts of the final subset dataset were used as input for CytoTRACE algorithm to infer directionality of differentiation (Gulati, Sikandar et al. 2020).

### RNA velocity

BAM files from each plate were processed using python command-line velocyto tool using run-smartseq2 command with GENCODE M21 genome and repeat masker annotation files, leading to a loom file for each plate containing spliced and unspliced transcript counts (La Manno, Soldatov et al. 2018). Using scvelo tool on python, genes having a shared count of spliced/unspliced of less than 20 were excluded and the 4000 top highly variable genes were kept (Bergen, Lange et al. 2020). PCA was performed on the spliced matrix, keeping 30 principal components and kNN neighbour graph was produced with k=15. Moments of spliced/unspliced abundances, velocity vectors and velocity graph were computed using default parameters. Extrapolated states were then projected on the UMAP embedding produced during the initial analysis.

### Terminal state identification using CellRank

CellRank was run to generate a transition matrix, using leiden clustering (resolution 1.5) and RNA velocity information weighted at 30% by connectivity generated from transcriptional information. To infer terminal states, the state space was decomposed into a set of 6 macrostates using GPCCA as estimator of the previously generated transition matrix. Among the 6 macrostates, 3 leiden clusters were selected as terminal states as one was indicating differentiated glial cells and two of these were indicative of two putative differentiated states of chromaffin cells. Absorption probabilities were generated for each of these terminal states. 15 cells for each tip of the terminal states were used for additional marker analysis, using t-test.

### Differentiation tree building using scFates

The CellRank results were converted to a principal tree using scFates python package. (*GitHub - LouisFaure/ScFates: A Scalable Python Suite for Tree Inference and Advanced Pseudotime Analysis from ScRNAseq Data*., n.d.) (Schumacher 2019). Briefly, a simplex embedding representing the forward terminal state probabilities was combined with the CytoTRACE values previously calculated. A 300 nodes principal graph was then inferred from this newly generated embedding, using SimplePPT method, which is based on the concept of a soft assignment matrix for each node to all cells (Zhang, Wang et al. 2015). This led to a tree composed of three branches capturing all three terminal states. The root of this tree was then automatically selected by first computing root cells via scvelo python package (tools.terminal_states, eps=0.01): For each nodes of the principal graph, the weighted average of the root cell probability was calculated using the soft assignment matrix previously generated. The node having the highest value of weighted averaged root cell was then selected as the root. Pseudotime values were then projected onto the cells along the trajectory, and all genes were tested for their association to the tree by using generalised additive model, comparing a full model (exp ∼ s(pseudotime)) to a reduced one (exp ∼ 1) via F-test. The test identified 6309 significant features after multiple hypothesis testing correction (FDR cut-off: 0.0001). Branching analysis of the two putative chromaffin terminal states was performed as following: First, **g**enes differentially upregulated between progenitor branch and terminal state were identified using the following linear model (exp ∼ pseudotime). Second, differentially expressed genes were then assigned between two post-bifurcation branches, using GAM by comparing a full model (exp ∼ s(pseudotime)*branch) to a reduced one (exp ∼ s(pseudotime)) via F-test.

### Emerging marker identification and correlation analysis

To identify early markers defining both glial and chromaffin fates, the inferred tree was subset by keeping only cells having a pseudotime up to the first detected ChC bifurcation. Then both trajectory toward glial and ChC were separately analysed, by test for associated genes along their respective path, with the same parameters as used for the whole tree. Both identified gene groups were then used to score the cells and assign them a fate using scanpy function score_genes. Cells were assigned to a fate if the score was higher than zero and higher than the other score. Cells not fulfilling these two criteria are defined as SCPs. Cells were manually selected in order to732 represent the different steps of differentiation. The local gene-gene correlation reflecting the coordination of early fate genes in SCPs was defined as a gene-gene Pearson correlation within these cells. The local correlation of a gene g with a module was assessed as a mean local correlation of that gene with the other genes comprising the module. Similarly, intra-module and inter-module correlations were taken to be the mean local gene-gene correlations of all possible gene pairs inside one module, or between the two modules, respectively (scFates, tl.slide_cors, default parameters).

### Embryo preparation for validations

For biological validations, adrenal glands were dissected from embryos E14.5, E16.5, E18.5 and P2 mice, fixed in 4% formaldehyde for 2-4 hours at +4°C, washed 3 times in PBS and transfered to 30% sucrose in 1xPBS. Cryoprotected tissue was embedded in OCT and stored at −80°C until cryosectioning took place. Samples were sectioned 12 μm thick and sections stored at −80°C until further analysis was performed.

### RNAscope® *in situ* hybridization and immunostaining

RNAscope *in situ* hybridization was performed according to manufacturer’s instructions using the RNAscope Multiplex Fluorescent Reagent Kit (ACD Biotechne) and the following commercially available probes: Mm-*Pnmt*-C3 (Cat No 426421-C3), Mm-*Lrp1b*-C1 (Cat No 491281), Mm-*Notum*-C1 (Cat.No: 428981), Mm-*Scnn1g*-C2 (Cat No 422091-C2), Mm-*Scg2*-C2 (Cat No 477691-C2), Mm-*Higd1c*-01-C3 (Cat No 527761-C3), Mm-*Casr*-C1 (Cat No 423451), Mm-*Myl1*-C2 (Cat No 548421-C2), Mm-*Rgs5*-C1 (Cat No 430181), Mm-*Caln1*-C1 (Cat No 551591). (Wang et al., 2012) Following RNAscope *in situ* hybridization immunofluorescence was performed by direct incubation with antibody at 4°C, overnight (rabbit polyclonal anti-TH, 1:1000, Pel-Freez Biologicals, #P40101-150). Following washes with 0.1% Tween in 1xPBS primary antibody was detected with secondary antibodies raised in donkey and conjugated with Alexa-488, 555, 647 (1:1000, Molecular Probes, Thermo Fisher Scientific).

### Imaging

Images were taken using Zeiss LSM800 confocal microscope equipped with 40x objective. Raw images were exported as TIFF files in ImageJ and figures were compiled using Adobe Photoshop and Illustrator.

### Statistics

Quantification of chromaffin subpopulations were obtained from adrenal glands from three embryos per developmental stage. Numbers of cells and mRNA transcripts were counted in ImageJ manually. Data were analyzed in GraphPad 8.1.1 (330) and presented as as mean± standard error of mean. No animals or data points were excluded from the analyses.

Proportion of Notum^+^/TH^+^ cells over total TH+ cells were 18.67±6.19 (E14.5, n=3), 35.59±4.83 (E16.5, n=3), 32.40±7.42 (E18.5, n=3), 71.34±28.66 (P2, n=3). Proportion of Higd1c^+^/TH^+^ cells over total TH+ cells were 96.0±4.0 (E14.5, n=3), 100±0 (E16.5, n=3), 49.13±9.86 (E18.5, n=3), 41.67±1.06 (P2, n=3). Proportion of Caln1^+^/TH^+^ cells over total TH+ cells were 17.09±3.79 (E14.5, n=3), 47.5±8.95 (E16.5, n=3), 21.04±19.20 (E18.5, n=3), 6.54±3.95 (P2, n=3). Proportion of Casr^+^/TH^+^ cells over total TH+ cells were 21.39±6.96 (E14.5, n=3), 44.59±6.41 (E16.5, n=3), 28.92±13.23 (E18.5, n=3), 29.38±4.64 (P2, n=3). Proportion of Pnmt^+^/TH^+^ cells over total TH+ cells were 35.08±6.13 (E14.5, n=3), 72.05±8.26 (E16.5, n=3), 100±0 (E18.5, n=3), 90.25±9.74 (P2, n=3). Proportion of Rgs5^+^/TH^+^ cells over total TH+ cells were 64.30±7.44 (E14.5, n=3), 50.99±23.89 (E16.5, n=3), 53.03±9.68 (E18.5, n=3), 49.41±10.80 (P2, n=3). Proportion of Myl1^+^/TH^+^ cells over total TH+ cells were 38.06±9.89 (E14.5, n=3), 43.38±16.36 (E16.5, n=3), 50.29±8.72 (E18.5, n=3), 5.99±4.76 (P2, n=3). Proportion of Scg2^+^/TH^+^ cells over total TH+ cells were 95.67±4.34 (E14.5, n=3), 100±0 (E16.5, n=3), 100±0 (E18.5, n=3), 14.29±1.38 (P2, n=3). Proportion of Lrp1b^+^/TH^+^ cells over total TH+ cells were 68.08±7.35 (E14.5, n=3), 100±0 (E16.5, n=3), 28.43±13.60 (E18.5, n=3), 76.09±23.92 (P2, n=3). Proportion of Scnn1g^+^/TH^+^ cells over total TH+ cells were 71.52±3.51 (E14.5, n=3), 8.66±1.65 (E16.5, n=3), 30.73±19.36 (E18.5, n=3), 11.17±2.93 (P2, n=3).

## Acknowledgements

We thank the Karolinska Eukaryotic Single Cell Genomics Facility (M. Erickson and K. Wallenborg) at SciLifeLab, Sweden for assistance with sequencing single cells.

## Author contribution

N.A. and P.A. acquired all biological data and performed the relevant analysis. L.F. performed computational analysis of single-cell data. M.E.K. and I.A. gave feedback on experimental aspects, supervised experimental approaches, and implemented the data interpretation. N.A., L.F., P.K., and M.E.K. made all figures containing data and resulting analysis. N.A., M.E.K, and I.A. designed the study, organized experimental work, and wrote the manuscript. All authors provided feedback on figures, manuscript composition, and structure.

## Data availability

The raw and processed data will be made available on the following GEO accession: GSE204700. To browse the data please use the link: https://adameykolab.srv.meduniwien.ac.at/ChC/.

The code for reproducing the bioinformatic analysis can be found on the following github repository: https://github.com/LouisFaure/ChromaffinFates_paper.

